# Inhibitor of the nuclear transport protein XPO1 enhances the anticancer efficacy of KRAS G12C inhibitors in preclinical models of KRAS G12C mutant cancers

**DOI:** 10.1101/2022.01.26.477874

**Authors:** Husain Yar Khan, Misako Nagasaka, Yiwei Li, Amro Aboukameel, Md. Hafiz Uddin, Rachel Sexton, Sahar Bannoura, Yousef Mzannar, Mohammed Najeeb Al-Hallak, Steve Kim, Rafic Beydoun, Yosef Landesman, Hirva Mamdani, Dipesh Uprety, Philip A. Philip, Ramzi M. Mohammad, Anthony F. Shields, Asfar S. Azmi

## Abstract

The identification of molecules that can bind covalently to KRAS G12C and lock it in an inactive GDP-bound conformation has opened the door to targeting KRAS G12C selectively. These agents have shown promise in preclinical tumor models and clinical trials. FDA has recently granted approval to sotorasib for KRAS G12C mutated non-small cell lung cancer (NSCLC). However, patients receiving these agents as monotherapy may not respond and generally develop drug resistance over time. This necessitates the development of multi-targeted approaches that can potentially sensitize tumors to KRAS inhibitors. We generated KRAS G12C inhibitor-resistant cell lines and observed that they exhibit sensitivity toward selinexor, a selective inhibitor of nuclear export protein exportin1 (XPO1), as a single agent. KRAS G12C inhibitor MRTX1257 in combination with selinexor suppressed the proliferation of KRAS G12C mutant cancer cell lines MiaPaCa-2 and NCI-H2122 in a synergistic manner. Moreover, combined treatment of selinexor with KRAS G12C inhibitors resulted in enhanced spheroid disintegration, reduction in the number and size of colonies formed by G12C mutant cancer cells. A combination of selinexor with KRAS G12C inhibitors potentiated the inhibition of KRAS expression in MiaPaCa-2 cells. NF-kB protein expression was also markedly reduced by selinexor and MRTX1257 combination. In an *in vivo* KRAS G12C cell-derived xenograft model, oral administration of a combination of selinexor and sotorasib was demonstrated to reduce tumor burden and enhance survival. In conclusion, we have shown that the nuclear transport protein XPO1 inhibitor can enhance the anticancer activity of KRAS G12C inhibitors in preclinical cancer models.

**Significance:** In this study, combining nuclear transport inhibitor selinexor with KRAS G12C inhibitors has resulted in potent antitumor effects in preclinical cancer models. This can be an effective combination therapy for cancer patients that do not respond or develop resistance to KRAS G12C inhibitor treatment.

## INTRODUCTION

RAS is one of the most frequently mutated oncogenes. In fact, more than 20% of all human cancers are associated with mutations in one of the three RAS isoforms, KRAS, HRAS, or NRAS [1, 2]. KRAS mutations are associated with a poor prognosis in general, notably in colorectal and pancreatic cancers. Mutations in KRAS are common in many solid tumors, most frequently occurring in 45% of colorectal, 35% of lung, and up to 90% of pancreatic cancers. In the United States alone, nearly 150,000 new cases of KRAS-mutated cancers are diagnosed each year across these three cancer types. More than half of all KRAS-driven cancers are caused by the three most common KRAS alleles, G12D, G12V, and G12C, which account for approximately 100,000 new cases in the US [3].

The KRAS gene encodes a small GTPase that acts as a molecular switch controlling key signaling pathways, such as the MAPK (RAF/MEK/ERK) and PI3K (PI3K/AKT/mTOR) pathways, which are responsible for cell proliferation and survival. KRAS protein alternate between the GDP-bound (inactive) and GTP-bound (active) states. GTP-bound KRAS stimulates the activation of many downstream signaling pathways. A key feature of oncogenic KRAS is impaired GTP hydrolysis, which results in increased flux through downstream pathways [4].

KRAS has long been considered undruggable. Due to its high affinity for GTP/GDP and the lack of a clear binding pocket, efforts to directly target KRAS have largely failed [5, 6]. However, a paradigm shift happened with the identification of a novel allosteric binding pocket under the switch II region of KRAS G12C protein that can be exploited for drug discovery. This led to the development of molecules that can covalently bind to G12C mutant KRAS at the cysteine 12 residue, thereby locking the protein in its inactive GDP-bound form that results in the inhibition of KRAS-dependent signaling, ultimately yielding antitumor activities [7-9]. This opened a window of opportunity to selectively target KRAS G12C protein using effective mutant-specific small molecule inhibitors, allowing KRAS to finally become druggable, albeit for a fraction of all KRAS-mutated tumors. Through this strategy several KRAS G12C inhibitors have been developed so far, including AMG510 (sotorasib) and MRTX849 (adagrasib). Both compounds have shown promising results in preclinical tumor models [10, 11] as well as in clinical trials [NCT03600883, NCT03785249]. Sotorasib has recently become the first KRAS G12C inhibitor to be granted accelerated approval by FDA as a second-line treatment for NSCLC patients carrying KRAS G12C mutation [12].

Although these KRAS G12C inhibitors have shown robust antitumor responses, but as targeted therapies, they are susceptible to the development of intrinsic or adaptive resistance, which can impede their prolonged therapeutic use [13,14]. There are already multiple indications that patients treated with these agents can develop drug resistance over time [15-18]. This necessitates the need for combination approaches that can potentially sensitize tumors to KRAS inhibitors when co-targeted.

The nuclear export protein exportin 1 (XPO1) plays a vital role in maintaining cellular homeostasis by mediating the export of a number of protein cargoes, including the majority of tumor suppressor proteins (TSPs), from the nucleus to the cytosol [19]. In many solid and hematological malignancies, increased expression of XPO1 has been observed which reportedly correlated with poor prognosis [20-21]. XPO1 overexpression enhances the export of TSPs to the cytosol, thereby preventing them from carrying out their normal function of cell growth regulation in the nucleus [22]. Therefore, XPO1 inhibition that can cause sequestering of TSPs within the nucleus, has emerged as an appealing anticancer strategy [19].

Interestingly, it has been reported that KRAS mutant NSCLC cells are dependent on XPO1 mediated nuclear export, rendering XPO1 a druggable vulnerability in KRAS mutant lung cancer [23]. Furthermore, the antitumor efficacy of XPO1 inhibitor selinexor against KRAS mutant lung cancer patient derived xenografts (PDXs) was recently demonstrated [24]. Moreover, XPO1 is linked to resistance to various standard-of-care chemotherapies and targeted therapies, which makes it a promising target for novel cancer therapies [25, 26]. In fact, XPO1 inhibitor selinexor (in combination with dexamethasone alone and with bortezomib and dexamethasone) has been approved for relapsed/refractory multiple myeloma patients [27] and as a monotherapy for patients with relapsed/refractory diffuse large B cell lymphoma [28].

In this study, we report for the first time that KRAS G12C inhibitor-resistant cell lines show sensitivity toward selinexor, providing a rationale for testing XPO1 inhibitor in combination with KRAS G12C inhibitors as an effective combination therapy. Using KRAS G12C mutant *in vitro* and *in vivo* preclinical models, we demonstrate enhanced anticancer activity of selinexor and KRAS G12C inhibitor combinations.

## MATERIALS AND METHODS

### Cell lines, drugs, and reagents

MiaPaCa-2 and NCI-H2122 cells were purchased from American Type Culture Collection (ATCC, Manassas, VA, USA). NCI “Rasless” mouse embryonic fibroblast (MEF) cell lines (KRAS 4B WT, G12C, G12D, G12V) were obtained from the National Cancer Institute (Rockville, MD, USA). MiaPaCa-2 and all the MEF cell lines were maintained in DMEM (Thermo Fisher Scientific, Waltham, MA, USA), while NCI-H2122 was maintained in RPMI1640 (Thermo Fisher Scientific, Waltham, MA, USA), supplemented with 10% fetal bovine serum (FBS), 100 U/mL penicillin, and 100 μg/mL streptomycin in a 5% CO2 atmosphere at 37 °C. The cell lines have been tested and authenticated in a core facility of the Applied Genomics Technology Center at Wayne State University. The method used for testing was short tandem repeat (STR) profiling using the PowerPlex® 16 System (Promega, Madison, WI, USA). MRTX1257 (ChemieTek, Indianapolis, IN, USA), AMG510, MRTX849 (Selleck Chemical LLC, Houston, TX) and selinexor (Karyopharm Therapeutics, Newton, MA, USA) were dissolved in DMSO to make 10 mM stock solutions. The drug control used for *in vitro* inhibitor experiments was cell culture media containing 0.1% DMSO.

### Generation of KRAS G12C inhibitor (AMG510 and MRTX1257) resistant cell lines

KRAS G12C mutant pancreatic ductal adenocarcinoma (PDAC) cell line, MiaPaCa-2, was maintained in long term cell culture exposed to incremental doses of AMG510 and MRTX1257 to develop drug resistance. MiaPaCa-2 cells, seeded at 60-70% confluence in DMEM and 10% FBS, were maintained in fresh drug containing medium changed every 3 days. The cells were passaged once they reach -90% confluence. The starting doses of the drugs were half of the IC_50_. Doses were doubled after every fifth passage of cell culture. The maximum dose the cells were exposed to was four times the IC50. After about 3 months (20 passages) of continuous drug exposure, the resulting pool of cells were collected and named as MIA-AMG-R and MIA-MRT-R. These cells were then treated with varying concentrations of the respective inhibitors and MTT assay was performed. Drug resistance was estimated by comparing the fold change in IC_50_s of the drug-primed and the unexposed parental cells.

### Cell viability assay and synergy analysis

Cells were seeded in 96-well culture plates at a density of 3 × 10^3^ cells per well. The growth medium was removed after overnight incubation and replaced with 100 μL of fresh medium containing the drug at various concentrations serially diluted from stock solution using OT-2 liquid handling robot (Opentrons, Queens, NY, USA). After 72 hours of exposure to the drug, MTT (3-(4,5-dimethylthiazol-2-yl)-2,5-diphenyltetrazolium bromide) assay was performed according to the procedure described previously [29]. Using the cell proliferation data (six replicates for each dose), IC_50_ values were calculated using the GraphPad Prism 4 software.

For the synergy analysis, cells were treated with three different concentrations of either MRTX1257/AMG510, or selinexor, or a combination of selinexor with MRTX1257/AMG510 at the corresponding doses for 72 hours (six replicates for each treatment). The drug proportion was kept constant across all the three dose combinations. Cell growth index was determined using MTT assay. The resulting cell growth data was used to generate isobolograms and calculate combination index (CI) values by the CalcuSyn software (Biosoft, Cambridge, UK).

### 3D culture and spheroid formation assay

MiaPaCa-2 and NCI-H2122 cells were trypsinized, collected as single cell suspensions using cell strainer and resuspended in sphere formation medium which was composed of 1:1 DMEM and F-12 nutrient mix supplemented with B-27 and N-2 (Thermo Fisher Scientific, Waltham, MA, USA). 1,000 cells were plated in each well of ultra-low attachment 6-well plates (Corning, Durham, NC, USA). Media was replenished every 3 days and spheroid growth was monitored. Spheroids growing in spheroid formation medium were exposed to either selinexor, or AMG510, or MRTX1257, or a combination of selinexor with either AMG510 or MRTX1257 twice a week for one week (three replicates for each treatment). At the end of the treatment, spheroids were counted under an inverted microscope and photographed.

### Colony formation assay

MiaPaCa-2 cells were seeded at a density of 500 cells per well in six well plates and exposed to single agent or combination drug treatments for 72 h. At the end of the treatment, drug containing media was removed and replaced with fresh media. The plates were incubated in the CO_2_ incubator for an additional ten days. After the incubation was over, media was removed from the wells of the plates and the colonies were fixed with methanol and stained with crystal violet for 15 minutes. The plates were then washed and dried before colonies were photographed. Colony number and sizes was later quantified by using NIH ImageJ 1.5Oi software.

### Preparation of total protein lysates and Western Blot analysis

1 × 10^6^ MiaPaCa-2 cells were grown in six 10 cm petri dishes overnight. The following day, cells were treated with the drugs as single agents or combinations for 12 hours. For total protein extraction, cells were lysed in RIPA buffer and protein concentrations were measured using BCA protein assay (PIERCE, Rockford, IL, USA). A total of 40 μg protein lysate from treated and untreated cells was resolved on a 10% SDS-PAGE and transferred onto a nitrocellulose membrane. The membrane was incubated with anti-KRAS (catalog # 517599; Santa Cruz Biotechnology, Santa Cruz, CA, USA), and anti-NF-κB p65 (catalog # 06-418; Millipore Sigma, Burlington, MA, USA) primary antibodies at 1:1000 dilution, while anti-β-actin (catalog # 47778; Santa Cruz Biotechnology, Santa Cruz, CA, USA) was used at a dilution of 1:3000. Incubation with HRP-linked secondary antibodies (catalog # 7074/7076; Cell Signaling, Danvers, MA, USA) was subsequently performed. The signal was detected using the ECL chemiluminescence detection system (Thermo Fisher Scientific, Waltham, MA, USA). Densitometric analysis of the data was performed using the ImageJ software (NIH, Bethesda, MD, USA).

### KRAS G12C cell-derived tumor xenograft study

*In vivo* studies were conducted under Wayne State University’s Institutional Animal Care and Use Committee (IACUC) approved protocol in accordance with the approved guidelines. Experiments were approved by the institute’s IACUC (Protocol # 18-12-0887). Post adaptation in our animal housing facility, 4-5 weeks old female ICR-SCID mice (Taconic Biosciences, Rensselaer, NY) were subcutaneously implanted with MiaPaCa-2 cells. 1×10^6^ cells suspended in 200 μL PBS were injected unilaterally into the left flank of donor mice using a BD 26Gx 5/8 1ml Sub-Q syringe. Once the tumors reached about 5-10% of the donor mice body weight, the donor mice were euthanized, tumors were harvested, and fragments were subsequently implanted into recipient mice. Seven days post transplantation, the recipient mice were randomly divided into four groups of 9 mice each and received either vehicle, or selinexor (15 mg/Kg once a week), or AMG510 (100 mg/Kg once daily), or their combination by oral gavage for 3 weeks. On completion of drug dosing, tumor tissue from control or treatment groups were used for RNA isolation and IHC analysis.

### RNA isolation and mRNA real-time RT-qPCR

Total RNAs from mouse tumors were extracted and purified using the RNeasy Mini Kit and RNase-free DNase Set (QIAGEN, Valencia, CA) following the protocol provided by the manufacturer. The expression levels of KRAS, XPO1, Erk2 and Bcl-2 in the mouse tumor tissues were analyzed by real-time RT-qPCR using High-Capacity cDNA Reverse Transcription Kit and SYBR Green Master Mixture from Applied Bio-systems (Waltham, MA, USA). The conditions and procedure for RT-qPCR have been described previously [29]. Sequences of primers used are listed in **Supplementary Table 1**.

### Immunostaining

Paraffin sections of the MiaPaCa-2 derived tumors were processed and stained with H&E and antibodies in a core facility at the Department of Oncology, Wayne State University. The following antibodies were used for immunohistochemistry staining: anti-Ki67 (catalog # M7240; Dako, Glostrup, Denmark) and anti-KRAS (catalog # 41-570-0; Fisher Scientific, Hampton, NH, USA) at 1:100 dilution, and anti-cleaved caspase-3 (catalog # 9664; Cell Signaling, Danvers, MA, USA) at 1:50 dilution.

### Statistical analysis

The student *t* test was used to compare statistically significant differences. Wherever suitable, the experiments were performed at least three times. The data were also subjected to unpaired two-tailed Student *t* test wherever appropriate, and *P < 0*.*05* was considered statistically significant.

### Data Availability

The data generated in this study are available upon request from the corresponding author.

## RESULTS

### Selinexor induces growth inhibition in PDAC cells resistant to KRAS G12C inhibitors

KRAS G12C inhibitor resistant cell lines were generated *in vitro* by continuous exposure of the KRAS G12C mutant PDAC cell line MiaPaCa-2 to increasing doses of AMG510 and MRTX1257. To establish the development of drug-resistance, we compared the IC_50_ values of the drug-exposed cell lines with the unexposed parental line. We observed 512-and 42-fold increase in the IC_50s_ of AMG510-resistant (MIA-AMG-R) and MRTX1257-resistant (MIA-MRT-R) MiaPaCa-2 cells, respectively, confirming that the cells had developed resistance to the respective KRAS G12C inhibitors (**Figure 1A**). Subsequently, both these drug-resistant cell lines (MIA-AMG-R and MIA-MRT-R) were treated with selinexor and were found to be sensitive to selinexor-induced cell growth inhibition (**Figure 1B**). This establishes that the KRAS G12C inhibitor-resistant cancer cells can potentially respond to selinexor.

**Figure 1:**
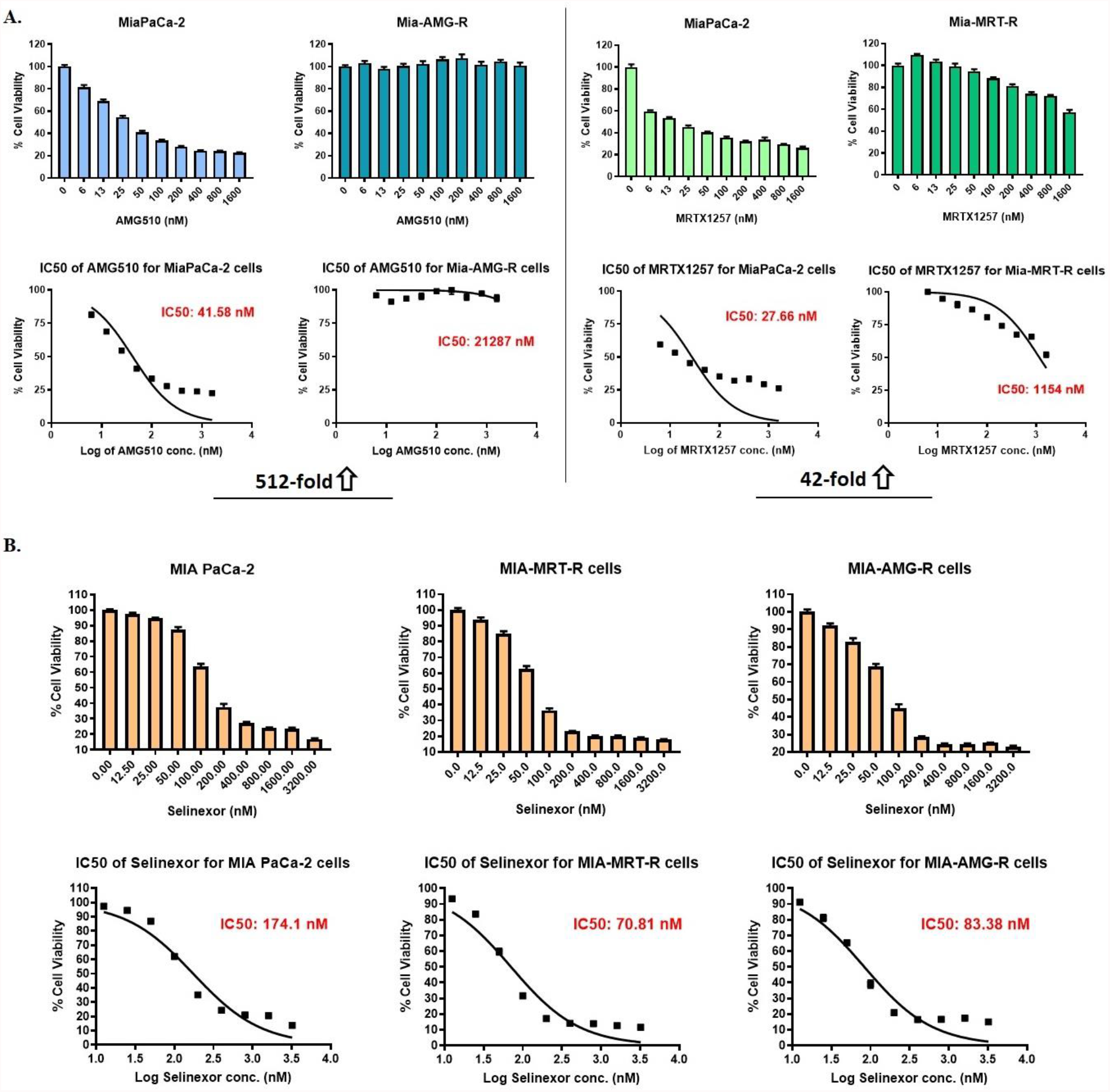
Selinexor induces growth inhibition in KRAS G12C inhibitor resistant cancer cells. (A) KRAS G12C mutant MiaPaCa-2 cells exposed to incremental doses of AMG510 and MRTX1257 in long term cell culture, eventually developed drug-resistance as shown by their unresponsiveness to drug treatment in MTT assay and several fold increase in the drug IC50 values compared to parental cells. (B) AMG510- and MRTX1257-resistant MiaPaCa-2 cell lines show sensitivity toward selinexor induced growth inhibition. Parental as well as resistant cells were treated with selinexor for 72h and MTT assay was performed as described in Methods. All results are expressed as percentage of control ± S.E.M of six replicates.

### Synthetic lethal interaction between XPO1 and KRAS

A computational systems approach called SLant (Synthetic Lethal analysis via Network topology) has recently been used for the prediction of human synthetic lethal (SSL) interactions via identifying and exploiting conserved patterns in protein interaction network topology [30]. We have obtained the experimentally validated synthetic lethal interactions of XPO1 using this approach from the Slorth database (http://slorth.biochem.sussex.ac.uk/welcome/index) and found interaction of XPO1 with KRAS to be synthetic lethal (**Supplementary Figure 1**).

### Combining selinexor with KRAS G12C inhibitors synergistically suppresses the proliferation of KRAS G12C mutant cells

KRAS G12C mutant NCI-H2122 (NSCLC) and MiaPaCa-2 (PDAC) cells were subjected to *in vitro* MRTX1257 and selinexor treatments at different dose combinations. As shown in **Figure 2A**, all three dose combinations tested demonstrated synergistic inhibition of NCI-H2122 cell proliferation (CI value < 1). For MiaPaCa-2 cells, synergistic effect (CI < 1) of the two drugs in suppressing cell growth was seen in at least two of the three combination doses tested (**Figure 2B**). AMG510 and selinexor combinations have more of an additive effect (CI equals 1) on the growth inhibition of NCI-H2122 and MiaPaCa-2 cells (**Supplementary Table 2**). These drug combinations were also tested on NCI “Rasless” MEFs carrying different KRAS mutations. Selinexor synergized with MRTX1257 at all dose combinations and with AMG510 at one of the higher dose combinations yielding suppressed growth of KRAS G12C mutant MEFs (**Supplementary Table 2**). As expected, the KRAS WT, KRAS G12D and KRAS G12V MEF cell lines were refractory to any such growth inhibition (**Supplementary Figure 2**).

**Figure 2:**
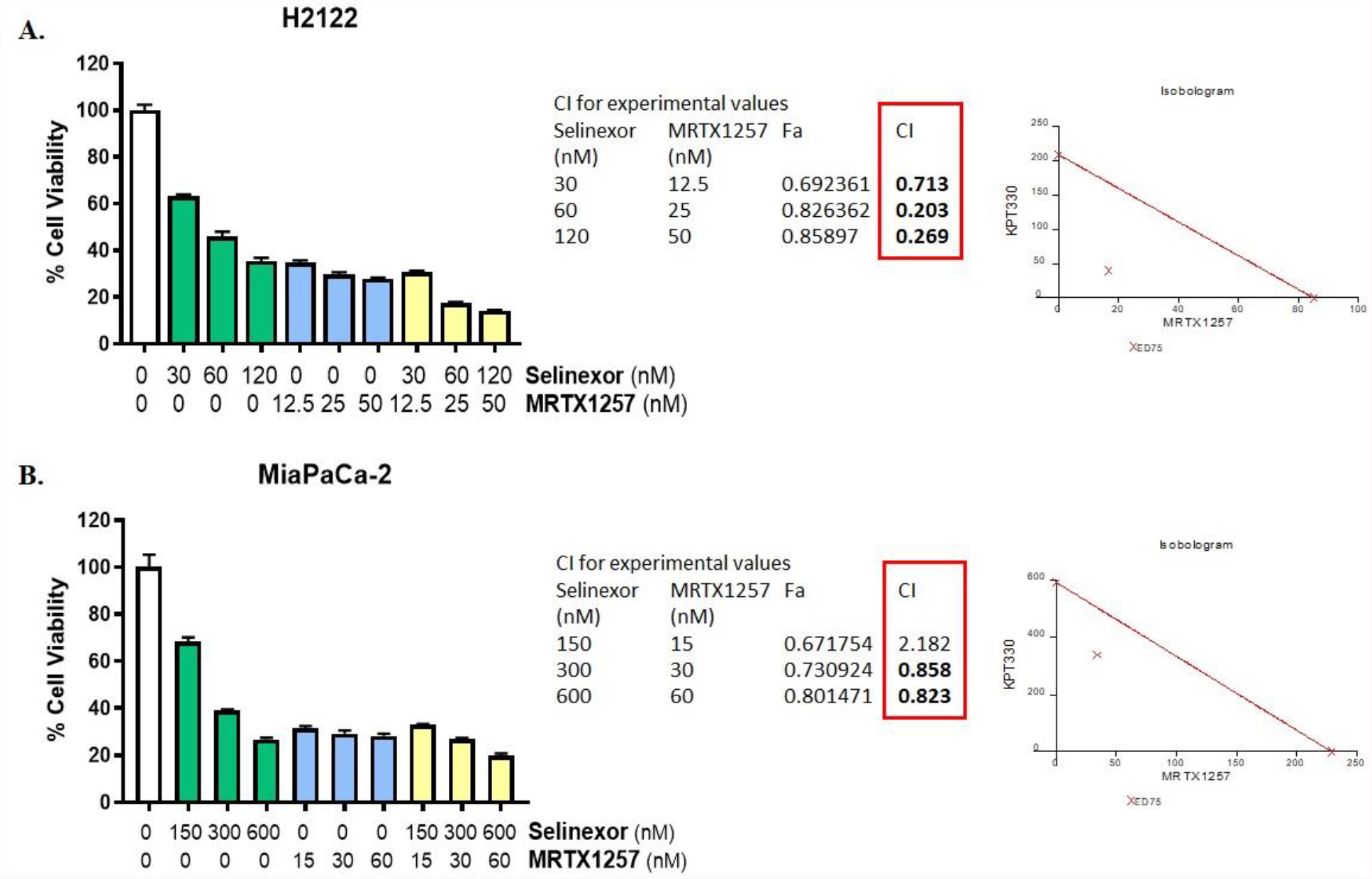
Selinexor and MRTX1257 show synergistic effects on the inhibition of cell proliferation *in vitro*. NCI-H2122 (A), and MiaPaCa-2 (B) cells were exposed to the indicated concentrations of either selinexor, MRTX1257, or a combination of both for 72 h, and cell proliferation was evaluated by MTT assay as described in Methods. CalcuSyn software was employed to generate isobolograms (shown on the right-hand side panel) and determine CI values from the resulting data. CI < 1 indicates synergistic effect of the drug combination at the corresponding doses. All results are expressed as percentage of control ± S.E.M of six replicates.

### Selinexor and KRAS G12C inhibitor combinations effectively disrupt the formation of KRAS G12C mutant cancer cell derived spheroids

Sensitivity of cells in 3D culture is considered to better predict *in vivo* efficacy, correlating well with drug response in xenograft models [31]. Therefore, we performed a spheroid formation assay, where combined treatment of selinexor with either MRTX1257 or AMG510 resulted in enhanced disruption of spheroids derived from MiaPaCa-2 and NCI-H2122 cell lines (**Figure 3**). Also, the total number of spheroids in the combination treated groups were significantly lower (p < 0.001) than that of the single agent selinexor treated group. Only with the MiaPaCa-2 derived spheroids, significant reduction (p < 0.01) in spheroid numbers of the AMG510 and selinexor combination compared to AMG510 alone was observed. These results demonstrate the efficacy of selinexor and AMG510 or MRTX1257 combinations in 3D cell growth models of KRAS G12C mutant PDAC and NSCLC.

**Figure 3:**
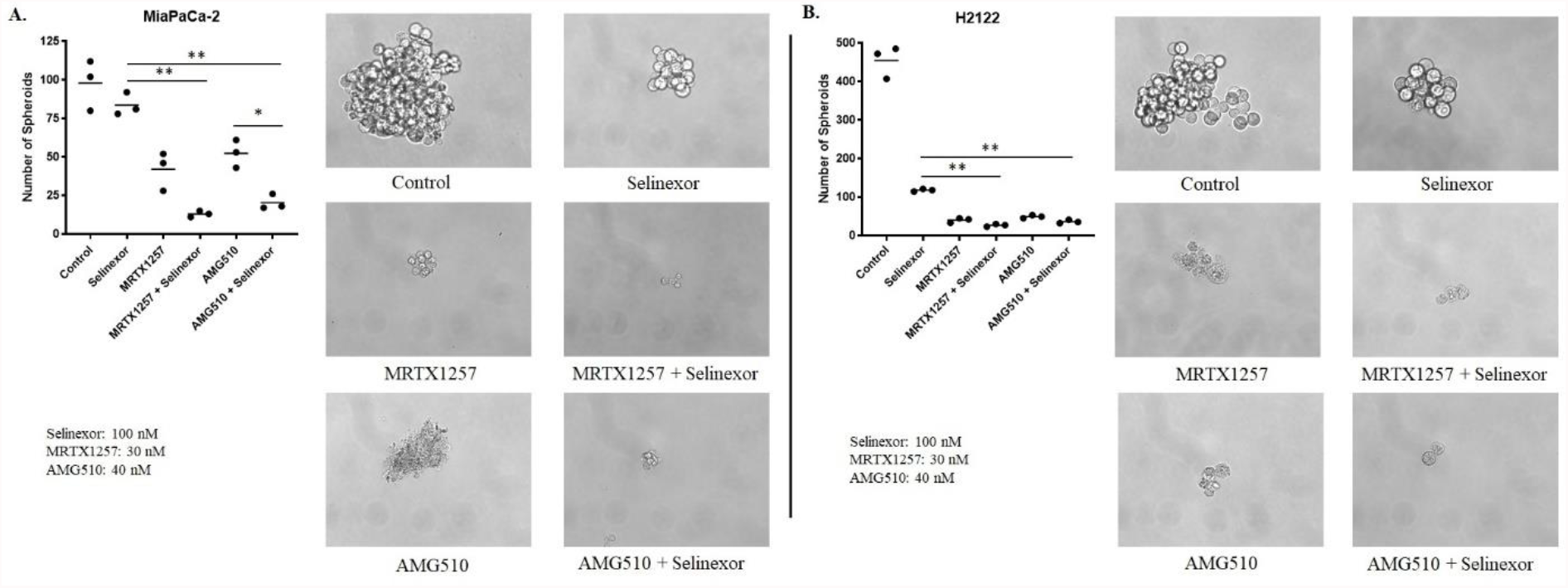
Selinexor in combination with MRTX1257 or AMG510 suppresses spheroid formation as well as significantly reduces the number of spheroids in 3D cultures of KRAS G12C mutant cancer cells. MiaPaCa-2 (A) and NCI-H2122 (B) cells were seeded in ultra-low attachment plates and treated with indicated concentrations of the drugs either as single agents or in combination for a week. Each treatment was performed in triplicate. At the end of treatment, spheroids were counted under the microscope and images were captured at 40x magnification.

### Combination of selinexor with KRAS G12C inhibitors reduces the clonogenic potential of KRAS G12C mutant cancer cells

The combinations of selinexor with MRTX1257, MRTX849 or AMG510 were evaluated for their effects on the colony formation ability of MiaPaCa-2 cells. Results of a clonogenic assay clearly demonstrate that the combination treatments of selinexor with each of the KRAS G12C inhibitors resulted in substantial decline in colony numbers as well as reduced average size of colonies formed by MiaPaCa-2 cells (**Figure 4**). Furthermore, this effect was more pronounced at the higher dose of KRAS G12C inhibitors tested (100 nM). These findings further underscore the efficacy of this combination approach in targeting KRAS G12C mutant cancer cells *in vitro*.

**Figure 4:**
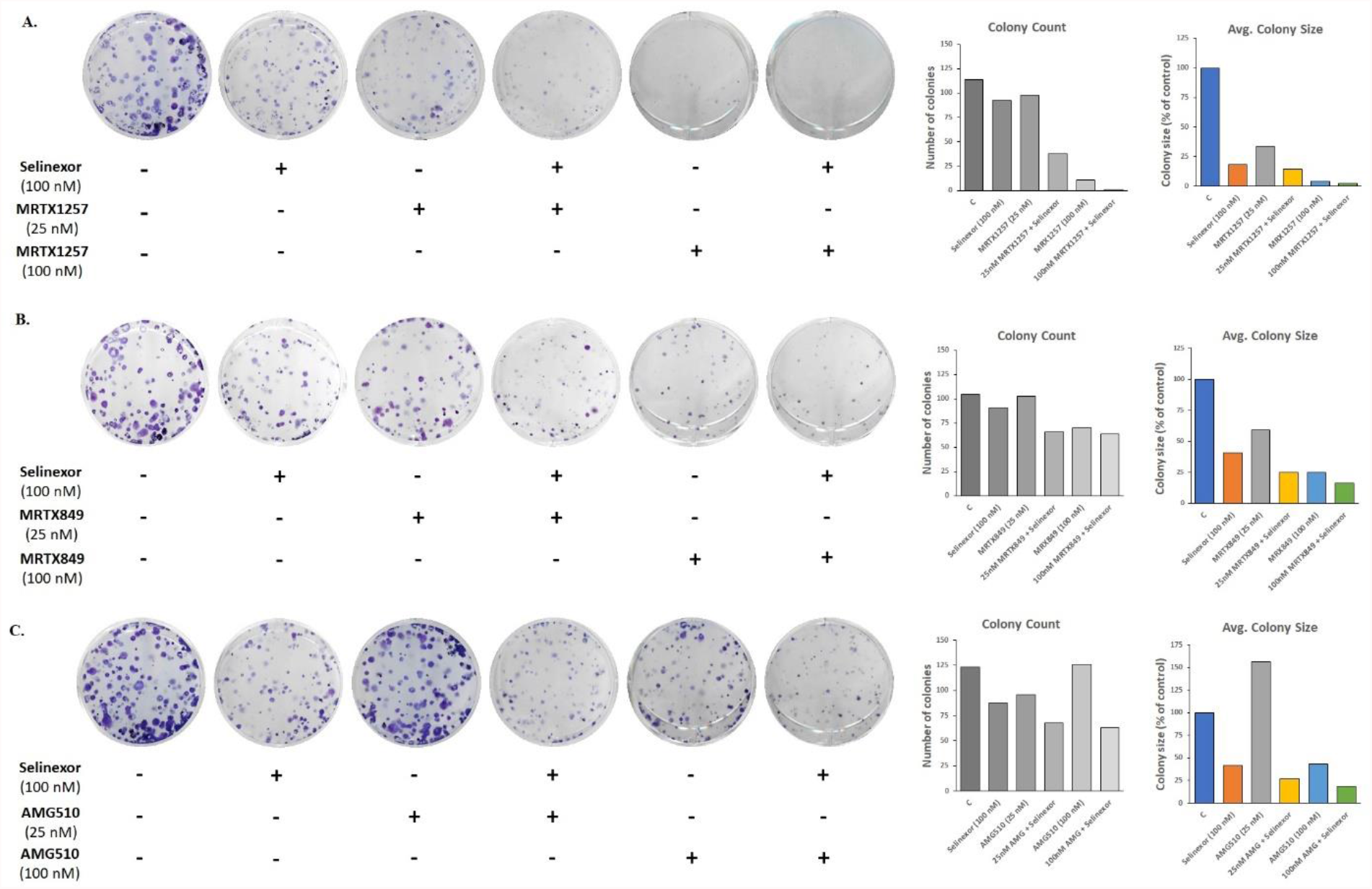
Combinations of selinexor with various KRAS G12C inhibitors inhibit the ability of KRAS^G12C^ mutant cancer cells to form colonies. MiaPaCa-2 cells were plated in 6-well plates (500 cells per well) and treated with combinations of selinexor with MRTX1257 (A), MRTX849 (B) and AMG510 (C) at the indicated concentrations for 72 h and colony formation assay was performed as described in Methods. Images of crystal violet-stained colonies were captured and NIH ImageJ 1.5Oi software was used to measure the number and size of colonies. Data is representative of three independent experiments.

### XPO1 and KRAS G12C inhibitor combination downregulates KRAS and NF-κB expression

As shown in **Figure 5**, the combination of selinexor with MRTX1257 or AMG510 potentiated the inhibition of KRAS expression in MiaPaCa-2 cells. NF-κB p65 protein expression was also markedly reduced in MiaPaCa-2 cells by selinexor and MRTX1257 combination compared to single agents. However, cells treated with selinexor and AMG510 combination showed only a slight decrease in the expression of NF-κB p65 in comparison to those treated with AMG510 alone. It was previously reported that the primary mechanism underlying XPO1 inhibitor sensitivity of KRAS-mutant lung cancer cell lines was intolerance to nuclear IκBα accumulation, with consequent inhibition of NF-κB signaling [28]. Although our data show a minor increase in NF-κB p65 subunit expression in MiaPaCa-2 cells exposed to single agent selinexor, the combination treatment nonetheless resulted in reduced NF-κB p65 expression, suggesting that the observed *in vitro* efficacy of the XPO1 inhibitor and KRAS G12C inhibitor combinations can be mechanistically attributed to the downregulation of NF-κB driven signaling.

**Figure 5:**
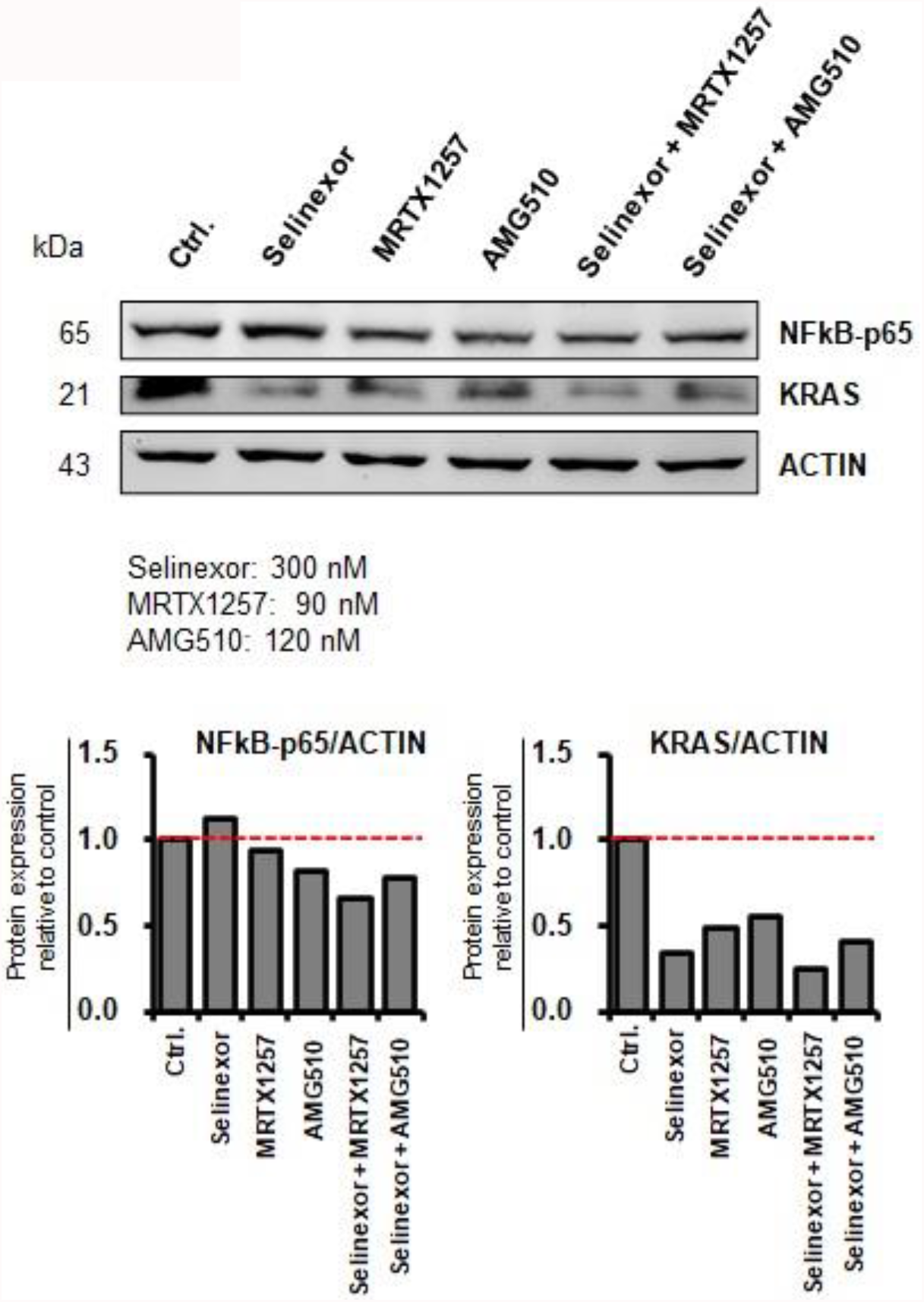
KRAS and NF-κB expression gets downregulated in cells treated with XPO1 and KRAS G12C inhibitor combination. MiaPaCa-2 cells were treated with indicated concentrations of either selinexor, or MRTX1257, or AMG510, or their combinations for 12 h. Protein extraction, determination of protein concentration, SDS-PAGE, and Western blot were performed as described in the Methods. β-actin was used as loading control. The quantitative analysis of mean pixel density of the blots was performed using NIH ImageJ 1.5Oi software.

### AMG510 and selinexor combination is more efficacious than single agent AMG510 in KRAS G12C mutant cell-derived xenograft model

In order to evaluate the *in vivo* effect of AMG510 either as a single agent or in combination with selinexor, a sub-cutaneous xenograft model of MiaPaCa-2 cells was established in ICR-SCID mice. The tumor-bearing mice were orally treated with selinexor (15 mg/kg; once a week), AMG510 (100 mg/Kg; daily) or the combination of AMG510 (100 mg/kg) and selinexor (15 mg/kg) for 3 weeks. Oral administration of AMG510 and selinexor combination showed greater tumor inhibition (**Figure 6A**) as well as enhanced survival of mice harboring MiaPaCa-2 subcutaneous xenografts (**Figure 6B**). Almost 22% of mice in the combination treatment group remained tumor free for as long as 150 days post tumor transplantation (**Figure 6B**). Additionally, the expression levels of KRAS, XPO1, ERK2 and Bcl-2 mRNA were found to be significantly decreased in the residual tumor samples from the combination group (**Figure 6C**). Further residual tumor profiling using IHC showed marked reduction in the proliferation marker, Ki67 and inhibition of KRAS in the combination group. In addition, the expression of pro-apoptotic marker, cleaved caspase-3 was high in the combination group (**Figure 6D**). These results, taken together, demonstrate the efficacy of AMG510 and selinexor combination *in vivo*.

**Figure 6:**
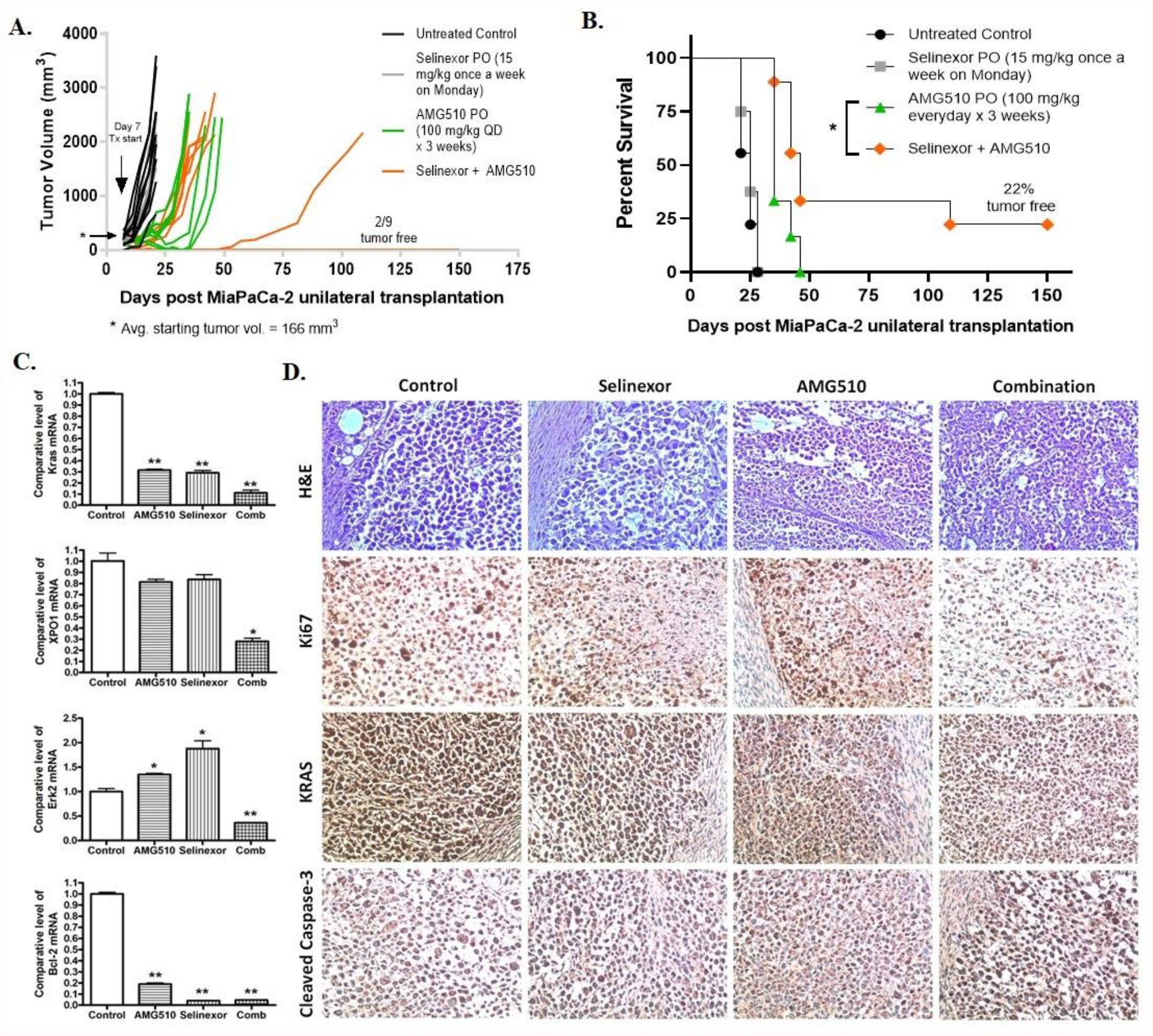
Preclinical antitumor efficacy of selinexor and KRAS G12C inhibitor combination in KRAS G12C CDX model. MiaPaCa-2 tumor xenografts were transplanted unilaterally in ICR-SCID mice, and the mice were randomly divided into four groups. Drug treatment was started one week after implanting xenografts when the average tumor volume reached 166 mm^3^. Selinexor was administered once a week at 15 mg/Kg, while AMG510 was given daily at 100 mg/Kg for 3 weeks. Tumor volume (A) and animal survival (B) were monitored up to 150 days post transplantation. Residual tumor tissues from each group were used for measuring mRNA levels of KRAS, XPO1, Erk2 and Bcl-2 (C), and performing immunohistochemical staining for Ki67, KRAS and cleaved caspase-3 (D).

## DISCUSSION

In this article, we show synergy between KRAS G12C inhibitors and nuclear protein export inhibitor, for the first time. Our combination approach of co-targeting KRAS G12C and XPO1 resulted in enhanced growth suppression of KRAS G12C mutant cells and cell-derived xenograft (CDX). This study brings forward a novel combination therapy for drug resistant KRAS G12C mutant tumors and provide preclinical rationale for the use of selinexor in a clinical setting to prevent or delay the development of resistance in patients receiving KRAS G12C inhibitor monotherapy.

Precision oncology has long sought to target KRAS oncoprotein directly. After decades of dismissing KRAS as untargetable, the development of inhibitors that can directly target KRAS G12C protein has rekindled hope. Researchers in the field have resumed their pursuit against KRAS with renewed vigor, especially after the early success of sotorasib and adagrasib in clinical trials [NCT03600883, NCT03785249]. The recent FDA approval of sotorasib has surely widened the panorama of treatment for patients harboring KRAS G12C mutation. However, preliminary clinical data and prior experience with other targeted therapies, such as EGFR and BRAF inhibitors, suggest that there are still many hurdles to overcome. Just like other targeted therapies, KRAS G12C inhibitors are anticipated to have limited efficacy as monotherapies and resistance develops in most patients, necessitating the use of combination therapies [4]. Hence, several combination approaches have been proposed, and some are currently undergoing clinical testing [11, 18, 32-34].

There is an increased realization for identifying KRAS associated synthetic lethality and developing small molecule inhibitors against such synthetic lethal targets. In a multi-genomic study, using 106 human NSCLC cell lines, Kim et al. [23] found that the nuclear transport machinery was selectively required for the survival of KRAS mutant cells that carry a broad range of phenotypic variation. The study further demonstrated that targeting nuclear export protein XPO1 with selinexor resulted in a robust synthetic lethal interaction with oncogenic KRAS both *in vitro* and *in vivo*. The identification of the existence of synthetic lethality between XPO1 and KRAS using the Slorth database in this study further rationalizes the significance of co-targeting XPO1 and KRAS.

In another study, selinexor treatment was found to effectively reduce tumor growth in ten KRAS mutant NSCLC PDXs irrespective of the type of KRAS mutation, indicating a general dependency of KRAS mutant cancers on XPO1 [24]. Moreover, XPO1 was identified to be a dependency in at least 90% of cancer cell lines in a genome wide CRISPR/Cas9 screen performed on 808 cell lines (Cancer Dependency Map Project), putting it into the category of ‘common essential gene’ [35]. Also, multiple reports implicate XPO1 to be a general vulnerability across several types of cancers [36-39].

Since XPO1 is overexpressed in a number of cancers [20, 21], it appears that XPO1 mediated nuclear export may be harnessed by various cancers as a general mechanism of oncogenesis. Therefore, a combination therapy involving XPO1 and KRAS G12C inhibitors can be a viable option, especially considering that preclinical and clinical studies have already reported the emergence of resistant subpopulations of cancer cells in response to KRAS G12C inhibitor monotherapy [13-16]. It can be speculated that these KRAS G12C inhibitor resistant cancer cells would be eradicated by the use of XPO1 inhibitor as a combination partner. This proposition was validated when we generated two KRAS G12C inhibitor-resistant cancer cell lines from KRAS G12C mutant parental cells (MiaPaCa-2) and found that both AMG510- and MRTX1257-resistant cell lines were indeed sensitive to the XPO1 inhibitor selinexor. Collectively, these findings imply that the inhibition of XPO1 activity could be a plausible therapeutic strategy for overcoming KRAS G12C resistance.

Our results demonstrate that the combinations of XPO1 inhibitor with KRAS G12C inhibitors can effectively inhibit the proliferation of KRAS G12C mutant cancer cells in 2D and 3D cultures. These combinations have been further shown to remarkably suppress the clonogenic potential of KRAS G12C mutant PDAC cells. In an *in vivo* KRAS G12C CDX model of PDAC, increased efficacy of selinexor and sotorasib combination in suppressing tumor growth and enhancing survival has been observed. These results support the *in vitro* finding that selinexor treatment can sensitize KRAS G12C inhibitor-resistant cancer cells. Further, this also suggests that combining selinexor with a KRAS G12C inhibitor sotorasib may have synergistic efficacy in cancer patients that have developed resistance to therapy with sotorasib (or other KRAS G12C inhibitors). In this regard, we have planned a Phase Ib/II study testing this combination in patients who have progressed on sotorasib. This novel combination therapy can potentially improve treatment outcomes in KRAS G12C mutant cancers.

## REFERENCES

1. AACR Project GENIE Consortium. AACR Project GENIE: Powering Precision Medicine through an International Consortium. Cancer Discov. 2017 Aug;7(8):818–831. doi: 10.1158/2159-8290.CD-17-0151. Epub 2017 Jun 1. PMID: 28572459; PMCID: PMC5611790.

2. Colicelli J. Human RAS superfamily proteins and related GTPases. Sci STKE. 2004 Sep 7;2004(250):RE13. doi: 10.1126/stke.2502004re13. PMID: 15367757; PMCID: PMC2828947.

3. Herdeis L, Gerlach D, McConnell DB, Kessler D. Stopping the beating heart of cancer: KRAS reviewed. Curr Opin Struct Biol. 2021 Jul 22;71:136–147. doi: 10.1016/j.sbi.2021.06.013. Epub ahead of print. PMID: 34303932.

4. Molina-Arcas M, Samani A, Downward J. Drugging the Undruggable: Advances on RAS Targeting in Cancer. Genes (Basel). 2021 Jun 10;12(6):899. doi: 10.3390/genes12060899. PMID: 34200676; PMCID: PMC8228461.

5. Matikas A, Mistriotis D, Georgoulias V, Kotsakis A. Targeting KRAS mutated non-small cell lung cancer: A history of failures and a future of hope for a diverse entity. Crit Rev Oncol Hematol. 2017 Feb;110:1–12. doi: 10.1016/j.critrevonc.2016.12.005. Epub 2016 Dec 9. PMID: 28109399.

6. Simanshu DK, Nissley DV, McCormick F. RAS Proteins and Their Regulators in Human Disease. Cell. 2017 Jun 29;170(1):17–33. doi: 10.1016/j.cell.2017.06.009. PMID: 28666118; PMCID: PMC5555610.

7. Janes MR, Zhang J, Li LS, Hansen R, Peters U, Guo X, Chen Y, Babbar A, Firdaus SJ, Darjania L, Feng J, Chen JH, Li S, Li S, Long YO, Thach C, Liu Y, Zarieh A, Ely T, Kucharski JM, Kessler LV, Wu T, Yu K, Wang Y, Yao Y, Deng X, Zarrinkar PP, Brehmer D, Dhanak D, Lorenzi MV, Hu-Lowe D, Patricelli MP, Ren P, Liu Y. Targeting KRAS Mutant Cancers with a Covalent G12C-Specific Inhibitor. Cell. 2018 Jan 25;172(3):578–589.e17. doi: 10.1016/j.cell.2018.01.006. PMID: 29373830.

8. Ostrem JM, Peters U, Sos ML, Wells JA, Shokat KM. K-Ras(G12C) inhibitors allosterically control GTP affinity and effector interactions. Nature. 2013 Nov 28;503(7477):548–51. doi: 10.1038/nature12796. Epub 2013 Nov 20. PMID: 24256730; PMCID: PMC4274051.

9. Patricelli MP, Janes MR, Li LS, Hansen R, Peters U, Kessler LV, Chen Y, Kucharski JM, Feng J, Ely T, Chen JH, Firdaus SJ, Babbar A, Ren P, Liu Y. Selective Inhibition of Oncogenic KRAS Output with Small Molecules Targeting the Inactive State. Cancer Discov. 2016 Mar;6(3):316–29. doi: 10.1158/2159-8290.CD-15-1105. Epub 2016 Jan 6. PMID: 26739882.

10. Canon J, Rex K, Saiki AY, Mohr C, Cooke K, Bagal D, Gaida K, Holt T, Knutson CG, Koppada N, Lanman BA, Werner J, Rapaport AS, San Miguel T, Ortiz R, Osgood T, Sun JR, Zhu X, McCarter JD, Volak LP, Houk BE, Fakih MG, O’Neil BH, Price TJ, Falchook GS, Desai J, Kuo J, Govindan R, Hong DS, Ouyang W, Henary H, Arvedson T, Cee VJ, Lipford JR. The clinical KRAS(G12C) inhibitor AMG 510 drives anti-tumour immunity. Nature. 2019 Nov;575(7781):217–223. doi: 10.1038/s41586-019-1694-1. Epub 2019 Oct 30. PMID: 31666701.

11. Hallin J, Engstrom LD, Hargis L, Calinisan A, Aranda R, Briere DM, Sudhakar N, Bowcut V, Baer BR, Ballard JA, Burkard MR, Fell JB, Fischer JP, Vigers GP, Xue Y, Gatto S, Fernandez-Banet J, Pavlicek A, Velastagui K, Chao RC, Barton J, Pierobon M, Baldelli E, Patricoin EF 3rd, Cassidy DP, Marx MA, Rybkin II, Johnson ML, Ou SI, Lito P, Papadopoulos KP, Jänne PA, Olson P, Christensen JG. The KRAS G12C Inhibitor MRTX849 Provides Insight toward Therapeutic Susceptibility of KRAS-Mutant Cancers in Mouse Models and Patients. Cancer Discov. 2020 Jan;10(1):54–71. doi: 10.1158/2159-8290.CD-19-1167. Epub 2019 Oct 28. PMID: 31658955; PMCID: PMC6954325.

12. Blair HA. Sotorasib: First Approval. Drugs. 2021 Sep;81(13):1573–1579. doi: 10.1007/s40265-021-01574-2. Erratum in: Drugs. 2021 Nov;81(16):1947. PMID: 34357500; PMCID: PMC8531079.

13. Xue JY, Zhao Y, Aronowitz J, Mai TT, Vides A, Qeriqi B, Kim D, Li C, de Stanchina E, Mazutis L, Risso D, Lito P. Rapid non-uniform adaptation to conformation-specific KRAS(G12C) inhibition. Nature. 2020 Jan;577(7790):421–425. doi: 10.1038/s41586-019-1884-x. Epub 2020 Jan 8. PMID: 31915379; PMCID: PMC7308074.

14. Adachi Y, Ito K, Hayashi Y, Kimura R, Tan TZ, Yamaguchi R, Ebi H. Epithelial-to-Mesenchymal Transition is a Cause of Both Intrinsic and Acquired Resistance to KRAS G12C Inhibitor in KRAS G12C-Mutant Non-Small Cell Lung Cancer. Clin Cancer Res. 2020 Nov 15;26(22):5962–5973. doi: 10.1158/1078-0432.CCR-20-2077. Epub 2020 Sep 8. PMID: 32900796.

15. Tanaka N, Lin JJ, Li C, Ryan MB, Zhang J, Kiedrowski LA, Michel AG, Syed MU, Fella KA, Sakhi M, Baiev I, Juric D, Gainor JF, Klempner SJ, Lennerz JK, Siravegna G, Bar-Peled L, Hata AN, Heist RS, Corcoran RB. Clinical Acquired Resistance to KRAS^G12C^ Inhibition through a Novel KRAS Switch-II Pocket Mutation and Polyclonal Alterations Converging on RAS-MAPK Reactivation. Cancer Discov. 2021 Aug;11(8):1913–1922. doi: 10.1158/2159-8290.CD-21-0365. Epub 2021 Apr 6. PMID: 33824136; PMCID: PMC8338755.

16. Awad MM, Liu S, Rybkin II, Arbour KC, Dilly J, Zhu VW, Johnson ML, Heist RS, Patil T, Riely GJ, Jacobson JO, Yang X, Persky NS, Root DE, Lowder KE, Feng H, Zhang SS, Haigis KM, Hung YP, Sholl LM, Wolpin BM, Wiese J, Christiansen J, Lee J, Schrock AB, Lim LP, Garg K, Li M, Engstrom LD, Waters L, Lawson JD, Olson P, Lito P, Ou SI, Christensen JG, Jänne PA, Aguirre AJ. Acquired Resistance to KRAS^G12C^ Inhibition in Cancer. N Engl J Med. 2021 Jun 24;384(25):2382–2393. doi: 10.1056/NEJMoa2105281. PMID: 34161704.

17. Molina-Arcas M, Moore C, Rana S, van Maldegem F, Mugarza E, Romero-Clavijo P, Herbert E, Horswell S, Li LS, Janes MR, Hancock DC, Downward J. Development of combination therapies to maximize the impact of KRAS-G12C inhibitors in lung cancer.Sci Transl Med. 2019 Sep 18;11(510):eaaw7999. doi: 10.1126/scitranslmed.aaw7999. PMID: 31534020; PMCID: PMC6764843.

18. Amodio V, Yaeger R, Arcella P, Cancelliere C, Lamba S, Lorenzato A, Arena S, Montone M, Mussolin B, Bian Y, Whaley A, Pinnelli M, Murciano-Goroff YR, Vakiani E, Valeri N, Liao WL, Bhalkikar A, Thyparambil S, Zhao HY, de Stanchina E, Marsoni S, Siena S, Bertotti A, Trusolino L, Li BT, Rosen N, Di Nicolantonio F, Bardelli A, Misale S. EGFR Blockade Reverts Resistance to KRAS^G12C^ Inhibition in Colorectal Cancer. Cancer Discov. 2020 Aug;10(8):1129–1139. doi: 10.1158/2159-8290.CD-20-0187. Epub 2020 May 19. PMID: 32430388; PMCID: PMC7416460.

19. Azmi AS, Uddin MH, Mohammad RM. The nuclear export protein XPO1 - from biology to targeted therapy. Nat Rev Clin Oncol. 2021 Mar;18(3):152–169. doi: 10.1038/s41571-020-00442-4. Epub 2020 Nov 10. Erratum in: Nat Rev Clin Oncol. 2020 Nov 24;: PMID: 33173198.

20. Birnbaum DJ, Finetti P, Birnbaum D, Mamessier E, Bertucci F. XPO1 Expression Is a Poor-Prognosis Marker in Pancreatic Adenocarcinoma. J Clin Med. 2019 Apr 30;8(5):596. doi: 10.3390/jcm8050596. PMID: 31052304; PMCID: PMC6572621.

21. van der Watt PJ, Maske CP, Hendricks DT, Parker MI, Denny L, Govender D, Birrer MJ, Leaner VD. The Karyopherin proteins, Crm1 and Karyopherin beta1, are overexpressed in cervical cancer and are critical for cancer cell survival and proliferation. Int J Cancer. 2009 Apr 15;124(8):1829–40. doi: 10.1002/ijc.24146. PMID: 19117056; PMCID: PMC6944291.

22. Nguyen KT, Holloway MP, Altura RA. The CRM1 nuclear export protein in normal development and disease. Int J Biochem Mol Biol. 2012;3(2):137–51. Epub 2012 May 18. PMID: 22773955; PMCID: PMC3388738.

23. Kim J, McMillan E, Kim HS, Venkateswaran N, Makkar G, Rodriguez-Canales J, Villalobos P, Neggers JE, Mendiratta S, Wei S, Landesman Y, Senapedis W, Baloglu E, Chow CB, Frink RE, Gao B, Roth M, Minna JD, Daelemans D, Wistuba II, Posner BA, Scaglioni PP, White MA. XPO1-dependent nuclear export is a druggable vulnerability in KRAS-mutant lung cancer. Nature. 2016 Oct 6;538(7623):114–117. doi: 10.1038/nature19771. Epub 2016 Sep 28. PMID: 27680702; PMCID: PMC5161658.

24. Rosen JC, Weiss J, Pham NA, Li Q, Martins-Filho SN, Wang Y, Tsao MS, Moghal N. Antitumor efficacy of XPO1 inhibitor Selinexor in KRAS-mutant lung adenocarcinoma patient-derived xenografts. Transl Oncol. 2021 Oct;14(10):101179. doi: 10.1016/j.tranon.2021.101179. Epub 2021 Jul 17. PMID: 34284202; PMCID: PMC8313753.

25. Seymour EK, Khan HY, Li Y, Chaker M, Muqbil I, Aboukameel A, Ramchandren R, Houde C, Sterbis G, Yang J, Bhutani D, Pregja S, Reichel K, Huddlestun A, Neveux C, Corona K, Landesman Y, Shah J, Kauffman M, Shacham S, Mohammad RM, Azmi AS, Zonder JA. Selinexor in Combination with R-CHOP for Frontline Treatment of Non-Hodgkin Lymphoma: Results of a Phase I Study. Clin Cancer Res. 2021 Jun 15;27(12):3307–3316. doi: 10.1158/1078-0432.CCR-20-4929. Epub 2021 Mar 30. PMID: 33785483; PMCID: PMC8197746.

26. Azmi AS, Khan HY, Muqbil I, Aboukameel A, Neggers JE, Daelemans D, Mahipal A, Dyson G, Kamgar M, Al-Hallak MN, Tesfaye A, Kim S, Shidham V, M Mohammad R, Philip PA. Preclinical Assessment with Clinical Validation of Selinexor with Gemcitabine and Nab-Paclitaxel for the Treatment of Pancreatic Ductal Adenocarcinoma. Clin Cancer Res. 2020 Mar 15;26(6):1338–1348. doi: 10.1158/1078-0432.CCR-19-1728. Epub 2019 Dec 12. PMID: 31831564; PMCID: PMC7073299.

27. Grosicki S, Simonova M, Spicka I, Pour L, Kriachok I, Gavriatopoulou M, Pylypenko H, Auner HW, Leleu X, Doronin V, Usenko G, Bahlis NJ, Hajek R, Benjamin R, Dolai TK, Sinha DK, Venner CP, Garg M, Gironella M, Jurczyszyn A, Robak P, Galli M, Wallington-Beddoe C, Radinoff A, Salogub G, Stevens DA, Basu S, Liberati AM, Quach H, Goranova-Marinova VS, Bila J, Katodritou E, Oliynyk H, Korenkova S, Kumar J, Jagannath S, Moreau P, Levy M, White D, Gatt ME, Facon T, Mateos MV, Cavo M, Reece D, Anderson LD Jr, Saint-Martin JR, Jeha J, Joshi AA, Chai Y, Li L, Peddagali V, Arazy M, Shah J, Shacham S, Kauffman MG, Dimopoulos MA, Richardson PG, Delimpasi S. Once-per-week selinexor, bortezomib, and dexamethasone versus twice-per-week bortezomib and dexamethasone in patients with multiple myeloma (BOSTON): a randomised, open-label, phase 3 trial. Lancet. 2020 Nov 14;396(10262):1563–1573. doi: 10.1016/S0140-6736(20)32292-3. PMID: 33189178.

28. Kalakonda N, Maerevoet M, Cavallo F, Follows G, Goy A, Vermaat JSP, Casasnovas O, Hamad N, Zijlstra JM, Bakhshi S, Bouabdallah R, Choquet S, Gurion R, Hill B, Jaeger U, Sancho JM, Schuster M, Thieblemont C, De la Cruz F, Egyed M, Mishra S, Offner F, Vassilakopoulos TP, Warzocha K, McCarthy D, Ma X, Corona K, Saint-Martin JR, Chang H, Landesman Y, Joshi A, Wang H, Shah J, Shacham S, Kauffman M, Van Den Neste E, Canales MA. Selinexor in patients with relapsed or refractory diffuse large B-cell lymphoma (SADAL): a single-arm, multinational, multicentre, open-label, phase 2 trial. Lancet Haematol. 2020 Jul;7(7):e511–e522. doi: 10.1016/S2352-3026(20)30120-4. PMID: 32589977.

29. Khan HY, Mpilla GB, Sexton R, Viswanadha S, Penmetsa KV, Aboukameel A, Diab M, Kamgar M, Al-Hallak MN, Szlaczky M, Tesfaye A, Kim S, Philip PA, Mohammad RM, Azmi AS. Calcium Release-Activated Calcium (CRAC) Channel Inhibition Suppresses Pancreatic Ductal Adenocarcinoma Cell Proliferation and Patient-Derived Tumor Growth. Cancers (Basel). 2020 Mar 22;12(3):750. doi: 10.3390/cancers12030750. PMID: 32235707; PMCID: PMC7140111.

30. Benstead-Hume G, Chen X, Hopkins SR, Lane KA, Downs JA, Pearl FMG. Predicting synthetic lethal interactions using conserved patterns in protein interaction networks. PLoS Comput Biol. 2019 Apr 17;15(4):e1006888. doi: 10.1371/journal.pcbi.1006888. PMID: 30995217; PMCID: PMC6488098.

31. Janes MR, Zhang J, Li LS, Hansen R, Peters U, Guo X, Chen Y, Babbar A, Firdaus SJ, Darjania L, Feng J, Chen JH, Li S, Li S, Long YO, Thach C, Liu Y, Zarieh A, Ely T, Kucharski JM, Kessler LV, Wu T, Yu K, Wang Y, Yao Y, Deng X, Zarrinkar PP, Brehmer D, Dhanak D, Lorenzi MV, Hu-Lowe D, Patricelli MP, Ren P, Liu Y. Targeting KRAS Mutant Cancers with a Covalent G12C-Specific Inhibitor. Cell. 2018 Jan 25;172(3):578–589.e17. doi: 10.1016/j.cell.2018.01.006. PMID: 29373830.

32. Fedele C, Li S, Teng KW, Foster CJR, Peng D, Ran H, Mita P, Geer MJ, Hattori T, Koide A, Wang Y, Tang KH, Leinwand J, Wang W, Diskin B, Deng J, Chen T, Dolgalev I, Ozerdem U, Miller G, Koide S, Wong KK, Neel BG. SHP2 inhibition diminishes KRASG12C cycling and promotes tumor microenvironment remodeling. J Exp Med. 2021 Jan 4;218(1):e20201414. doi: 10.1084/jem.20201414. PMID: 33045063; PMCID: PMC7549316.

33. Brown WS, McDonald PC, Nemirovsky O, Awrey S, Chafe SC, Schaeffer DF, Li J, Renouf DJ, Stanger BZ, Dedhar S. Overcoming Adaptive Resistance to KRAS and MEK Inhibitors by Co-targeting mTORC1/2 Complexes in Pancreatic Cancer. Cell Rep Med. 2020 Nov 17;1(8):100131. doi: 10.1016/j.xcrm.2020.100131. PMID: 33294856; PMCID: PMC7691443.

34. Zhang B, Zhang Y, Zhang J, Liu P, Jiao B, Wang Z, Ren R. Focal Adhesion Kinase (FAK) Inhibition Synergizes with KRAS G12C Inhibitors in Treating Cancer through the Regulation of the FAK-YAP Signaling. Adv Sci (Weinh). 2021 Aug;8(16):e2100250. doi: 10.1002/advs.202100250. Epub 2021 Jun 20. PMID: 34151545; PMCID: PMC8373085.

35. Meyers RM, Bryan JG, McFarland JM, Weir BA, Sizemore AE, Xu H, Dharia NV, Montgomery PG, Cowley GS, Pantel S, Goodale A, Lee Y, Ali LD, Jiang G, Lubonja R, Harrington WF, Strickland M, Wu T, Hawes DC, Zhivich VA, Wyatt MR, Kalani Z, Chang JJ, Okamoto M, Stegmaier K, Golub TR, Boehm JS, Vazquez F, Root DE, Hahn WC, Tsherniak A. Computational correction of copy number effect improves specificity of CRISPR-Cas9 essentiality screens in cancer cells. Nat Genet. 2017 Dec;49(12):1779–1784. doi: 10.1038/ng.3984. Epub 2017 Oct 30. PMID: 29083409; PMCID: PMC5709193.

36. Sexton R, Mahdi Z, Chaudhury R, Beydoun R, Aboukameel A, Khan HY, Baloglu E, Senapedis W, Landesman Y, Tesfaye A, Kim S, Philip PA, Azmi AS. Targeting Nuclear Exporter Protein XPO1/CRM1 in Gastric Cancer. Int J Mol Sci. 2019 Sep 28;20(19):4826. doi: 10.3390/ijms20194826. PMID: 31569391; PMCID: PMC6801932.

37. Chen Y, Camacho SC, Silvers TR, Razak AR, Gabrail NY, Gerecitano JF, Kalir E, Pereira E, Evans BR, Ramus SJ, Huang F, Priedigkeit N, Rodriguez E, Donovan M, Khan F, Kalir T, Sebra R, Uzilov A, Chen R, Sinha R, Halpert R, Billaud JN, Shacham S, McCauley D, Landesman Y, Rashal T, Kauffman M, Mirza MR, Mau-Sørensen M, Dottino P, Martignetti JA. Inhibition of the Nuclear Export Receptor XPO1 as a Therapeutic Target for Platinum-Resistant Ovarian Cancer. Clin Cancer Res. 2017 Mar 15;23(6):1552–1563. doi: 10.1158/1078-0432.CCR-16-1333. Epub 2016 Sep 20. PMID: 27649553.

38. Arango NP, Yuca E, Zhao M, Evans KW, Scott S, Kim C, Gonzalez-Angulo AM, Janku F, Ueno NT, Tripathy D, Akcakanat A, Naing A, Meric-Bernstam F. Selinexor (KPT-330) demonstrates anti-tumor efficacy in preclinical models of triple-negative breast cancer. Breast Cancer Res. 2017 Aug 15;19(1):93. doi: 10.1186/s13058-017-0878-6. PMID: 28810913; PMCID: PMC5557476.

39. Sun H, Hattori N, Chien W, Sun Q, Sudo M, E-Ling GL, Ding L, Lim SL, Shacham S, Kauffman M, Nakamaki T, Koeffler HP. KPT-330 has antitumour activity against non-small cell lung cancer. Br J Cancer. 2014 Jul 15;111(2):281–91. doi: 10.1038/bjc.2014.260. Epub 2014 Jun 19. PMID: 24946002; PMCID: PMC4102938.

