## Supplemental Data for "Inhibitor of the nuclear transport protein XPO1 enhances the anticancer efficacy of KRAS G12C inhibitors in preclinical models of KRAS G12C mutant cancers"

### Slide 1
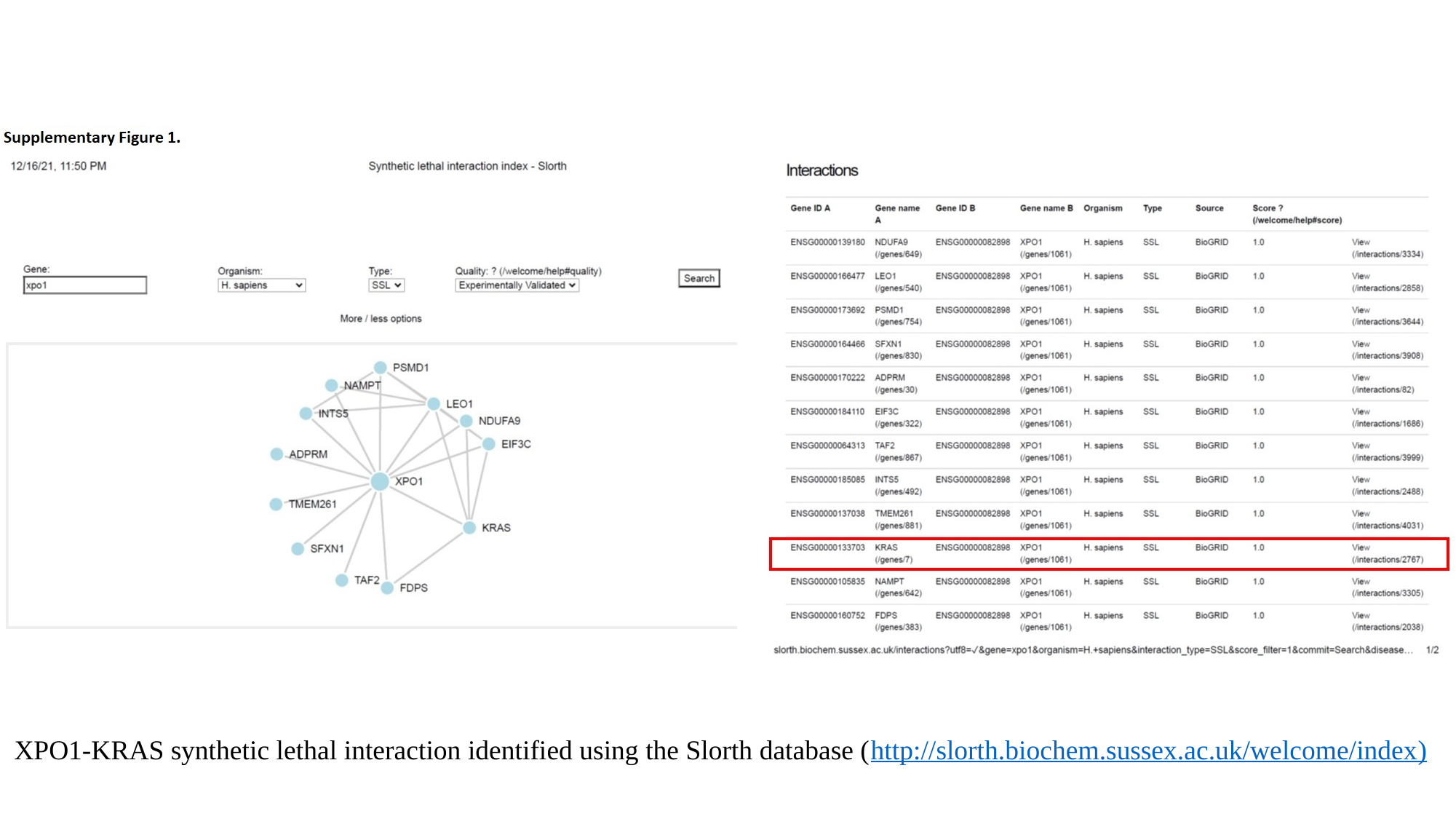

XPO1-KRAS synthetic lethal interaction identified using the Slorth database (http://slorth.biochem.sussex.ac.uk/welcome/index)

### Slide 2
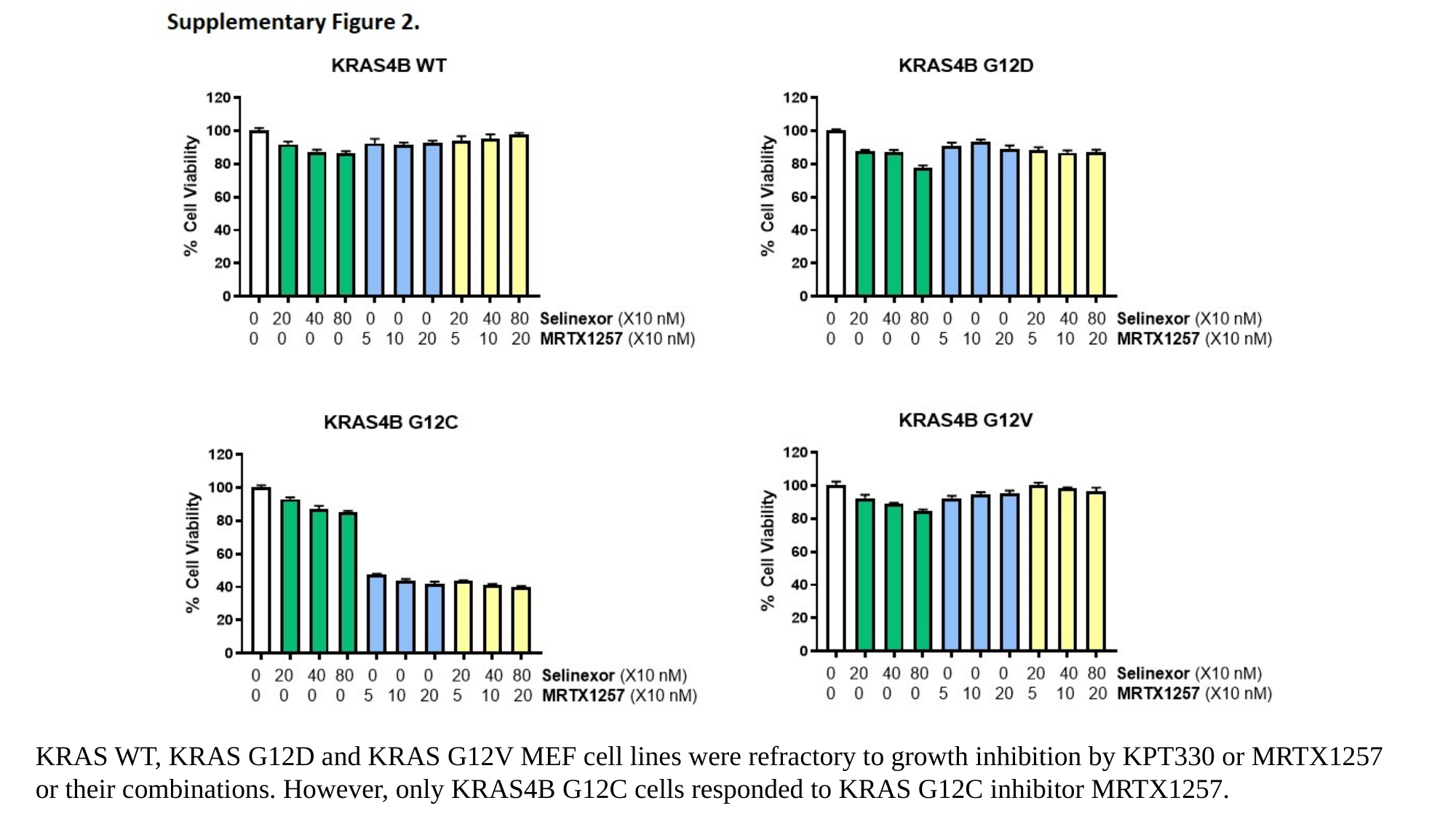

KRAS WT, KRAS G12D and KRAS G12V MEF cell lines were refractory to growth inhibition by KPT330 or MRTX1257 or their combinations. However, only KRAS4B G12C cells responded to KRAS G12C inhibitor MRTX1257.

### Slide 3
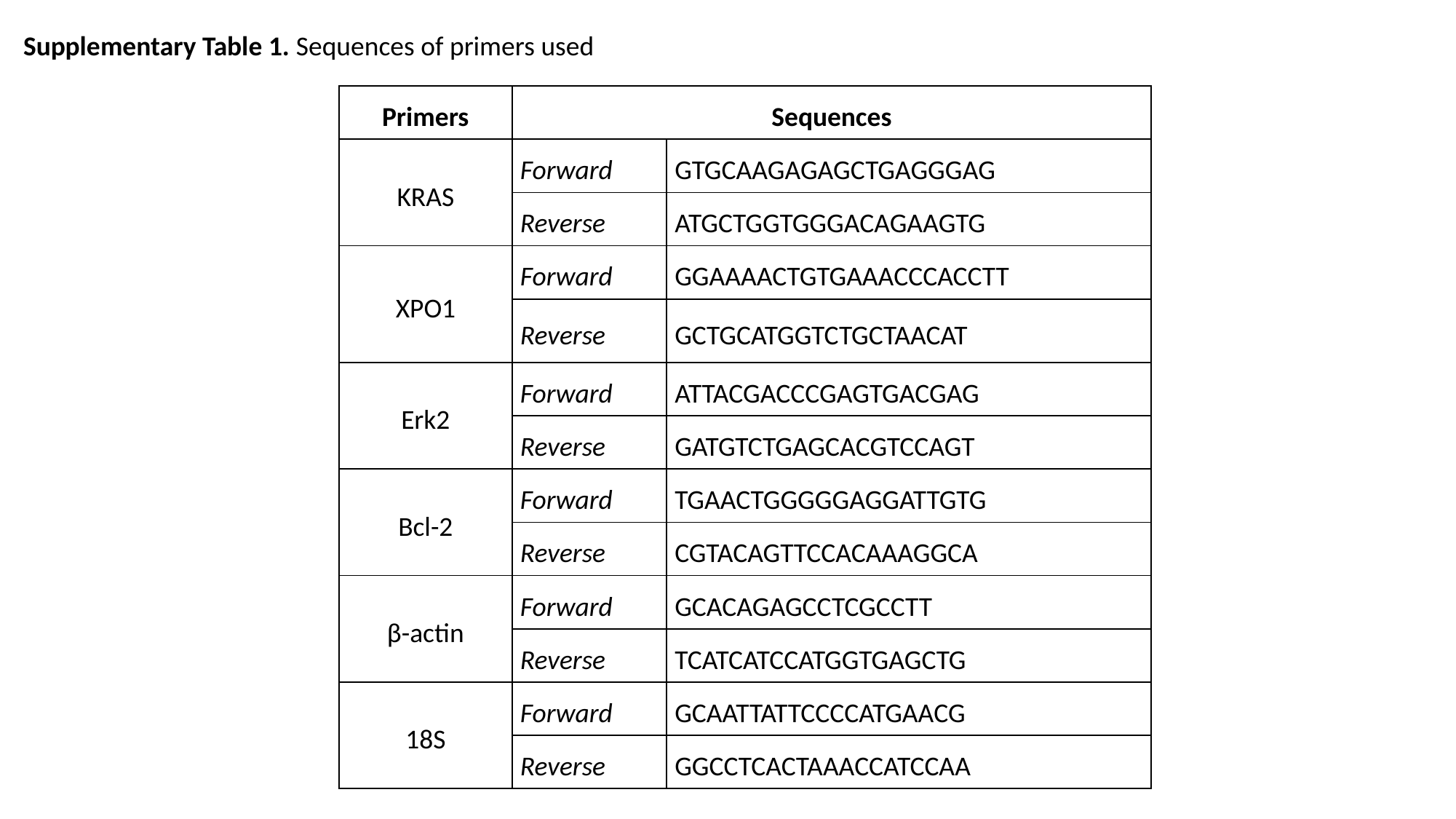

Supplementary Table 1. Sequences of primers used
| Primers | Sequences | |
| --- | --- | --- |
| KRAS | Forward | GTGCAAGAGAGCTGAGGGAG |
| | Reverse | ATGCTGGTGGGACAGAAGTG |
| XPO1 | Forward | GGAAAACTGTGAAACCCACCTT |
| | Reverse | GCTGCATGGTCTGCTAACAT |
| Erk2 | Forward | ATTACGACCCGAGTGACGAG |
| | Reverse | GATGTCTGAGCACGTCCAGT |
| Bcl-2 | Forward | TGAACTGGGGGAGGATTGTG |
| | Reverse | CGTACAGTTCCACAAAGGCA |
| β-actin | Forward | GCACAGAGCCTCGCCTT |
| | Reverse | TCATCATCCATGGTGAGCTG |
| 18S | Forward | GCAATTATTCCCCATGAACG |
| | Reverse | GGCCTCACTAAACCATCCAA |

### Slide 4
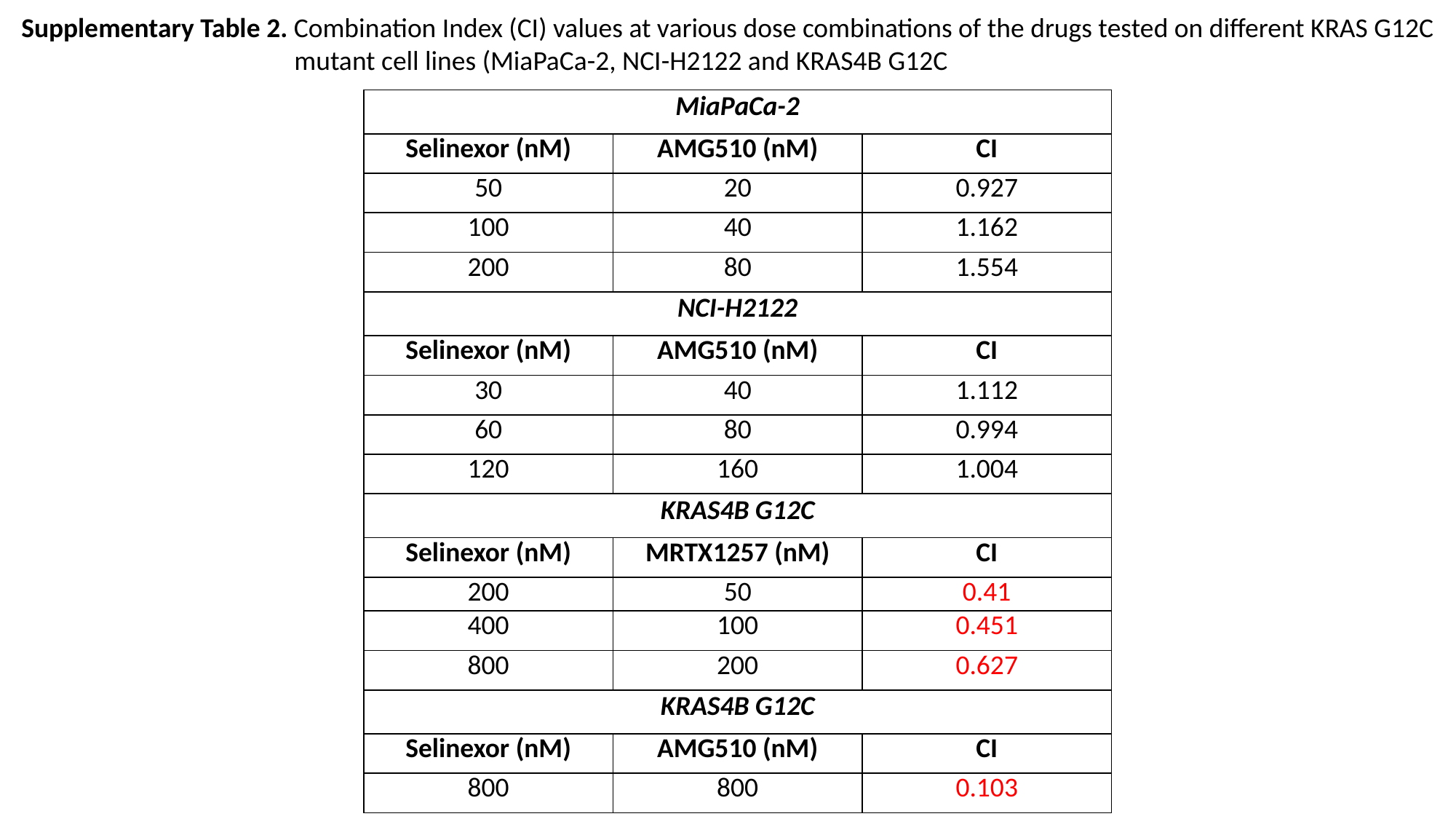

Supplementary Table 2. Combination Index (CI) values at various dose combinations of the drugs tested on different KRAS G12C mutant cell lines (MiaPaCa-2, NCI-H2122 and KRAS4B G12C
| MiaPaCa-2 | AMG510 (nM) | CI |
| --- | --- | --- |
| Selinexor (nM) | AMG510 (nM) | CI |
| 50 | 20 | 0.927 |
| 100 | 40 | 1.162 |
| 200 | 80 | 1.554 |
| NCI-H2122 | | |
| Selinexor (nM) | AMG510 (nM) | CI |
| 30 | 40 | 1.112 |
| 60 | 80 | 0.994 |
| 120 | 160 | 1.004 |
| KRAS4B G12C | | |
| Selinexor (nM) | MRTX1257 (nM) | CI |
| 200 | 50 | 0.41 |
| 400 | 100 | 0.451 |
| 800 | 200 | 0.627 |
| KRAS4B G12C | | |
| Selinexor (nM) | AMG510 (nM) | CI |
| 800 | 800 | 0.103 |
